# Structural and functional synaptic plasticity induced by convergent synapse loss requires co-innervation in the Drosophila neuromuscular circuit

**DOI:** 10.1101/2020.06.12.147421

**Authors:** Yupu Wang, Meike Lobb-Rabe, James Ashley, Robert A. Carrillo

**Affiliations:** Department of Molecular Genetics and Cellular Biology, University of Chicago, Chicago, IL 60637; Grossman Institute for Neuroscience, Quantitative Biology and Human Behavior, University of Chicago, Chicago, IL 60637; Committe on Development, Regeneration, and Stem Cell Biology, University of Chicago, Chicago, IL 60637; Program in Cell and Molecular Biology, University of Chicago, Chicago, IL 60637

## Abstract

Throughout the nervous system, the convergence of two or more presynaptic inputs on a target cell is commonly observed. The question we ask here is to what extent converging inputs influence each other’s structural and functional synaptic plasticity. In complex circuits, isolating individual inputs is difficult because postsynaptic cells can receive thousands of inputs. An ideal model to address this question is the *Drosophila* larval neuromuscular junction where each postsynaptic muscle cell receives inputs from two glutamatergic types of motor neurons (MNs), known as 1b and 1s MNs. Notably, each muscle is unique and receives input from a different combination of 1b and 1s motor neurons. We surveyed synapses on multiple muscles for this reason. Here, we identified a cell-specific promoter to ablate 1s MNs after innervation. Additionally, we genetically blocked 1s innervation. Then we measured 1b MN structural and functional responses using electrophysiology and microscopy. For all muscles, 1s MN ablation resulted in 1b MN synaptic expansion and increased basal neurotransmitter release. This demonstrates that 1b MNs can compensate for the loss of convergent inputs. However, only a subset of 1b MNs showed compensatory evoked activity, suggesting spontaneous and evoked plasticity are independently regulated. Finally, we used *DIP-α* mutants that block 1s MN synaptic contacts; this eliminated robust 1b synaptic plasticity, raising the possibility that muscle co-innervation may define an activity “set point” that is referenced when subsequent synaptic perturbations occur. This model can be tested in more complex circuits to determine if co-innervation is fundamental for input-specific plasticity.

**SIGNIFICANCE STATEMENT:** In complex neural circuits, multiple converging inputs contribute to the output of each target cell. Thus, each input must be regulated, but whether adjacent inputs contribute to this regulation is unclear. To examine input-specific synaptic plasticity in a structurally and functionally tractable system, we turn to the *Drosophila* neuromuscular circuit. Each muscle is innervated by a unique pair of motor neurons. Removal of one neuron after innervation causes the adjacent neuron to increase synaptic outgrowth and functional output. However, this is not a general feature since each MN differentially compensates. Also, robust compensation requires co-innervation by both neurons. Understanding how neurons respond to perturbations in adjacent neurons will provide insight into nervous system plasticity in both healthy and diseased states.

## INTRODUCTION

The nervous system is characterized by complex wiring patterns that include different neurons converging onto the same postsynaptic cell. This wiring paradigm is found in pyramidal neurons that receive input from excitatory and inhibitory contacts (Megias et al., 2001), and in esophageal striated muscles that receive enteric and vagal nerve inputs (Kallmunzer et al., 2008; Neuhuber and Worl, 2016). While dynamic regulation of individual synapses has been examined (Berry and Nedivi, 2017; Kruijssen and Wierenga, 2019), the interplay between adjacent synapses from different neurons has been predominately studied by monitoring postsynaptic spine changes in response to perturbations of adjacent synapses (Chistiakova et al., 2019; Hedrick et al., 2016; Jungenitz et al., 2018; Oh et al., 2015). Due to the complex and dense nature of these circuits, it is difficult to study the changes induced by the absence of a single presynaptic contact or an entire presynaptic neuron with several contacts. Understanding how neurons respond to these perturbations will shed light on the etiology of neurodegenerative disorders, such as traumatic brain injury (TBI) and amyotrophic lateral sclerosis (ALS), which display progressive neuronal cell death and devastating functional consequences (Chauhan, 2014; Cluskey and Ramsden, 2001).

Analysis of individual inputs in the central nervous system is complicated by the high density of converging inputs on the same cell. The *Drosophila* larval neuromuscular circuit, however, circumvents this with a simple, hard-wired connectivity map (Figure 1A). The body plan is segmentally repeated and bilateral symmetry allows division into hemisegments, which are comprised of 35 motor neurons (MNs) and 30 postsynaptic muscles. Like most vertebrate central synapses, the individual glutamatergic neuromuscular contacts consist of axon terminal swellings, called boutons, and elaborate postsynaptic complexes (Collins and DiAntonio, 2007; Menon et al., 2013; Van Vactor and Sigrist, 2017). Each bouton houses several active zones (AZs) that enable neurotransmission. Larval muscles are innervated by two main MNs (named for anatomical observations of morphology), type 1b (big) and 1s (small), which resemble tonic and phasic neurons, respectively (Schaefer et al., 2010). Most 1b MNs innervate single muscles, whereas 1s MNs innervate functional muscle groups (Hoang and Chiba, 2001; Landgraf et al., 1997; Lnenicka and Keshishian, 2000), highlighting important structural and functional distinctions between these classes of neurons.

**Figure 1.**
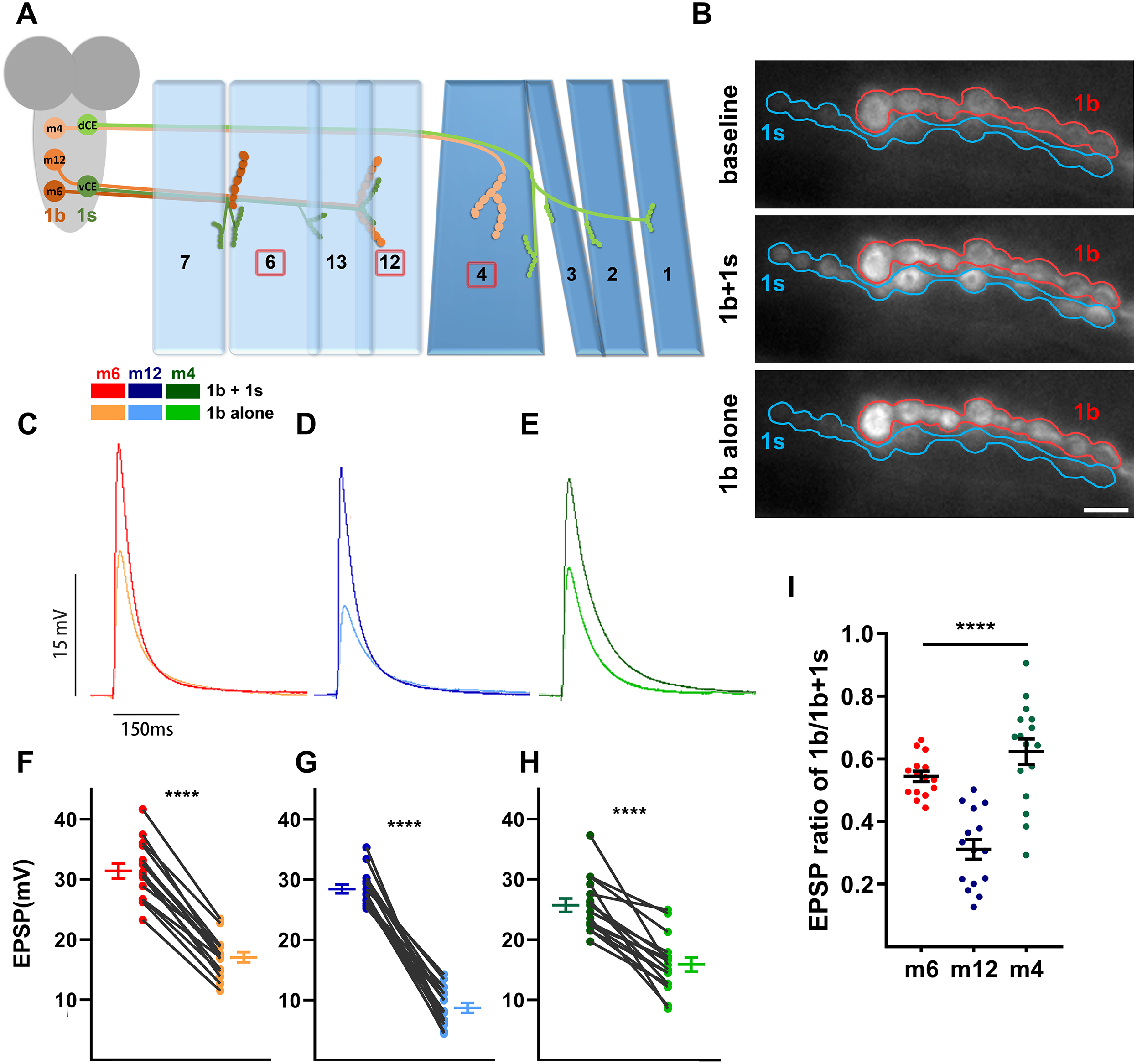
1b and 1s inputs differentially contribute to the total EPSPs. (**A**) Schematic of the innervation pattern of a subset of 1s motor neurons (vCE: dark green, dCE: light green) and 1b motor neurons (m6-1b: rust, m12-1b: orange, m4-1b: peach). Muscles analyzed in this study are marked by the red boxes. (**B**) Representative frames showing the baseline fluorescence (top), 1b+1s firing event (middle) and 1b alone firing event (bottom), of m6 in MHC-CD8-SynapGCaMP6f larvae (1b: red, 1s: blue) (also see Movie 1). (**C-E**) Representative traces of 1b+1s and 1b alone on (**C**) m6, (**D**) m12, and (**E**) m4. (**F-H**) Paired EPSP amplitudes from (**F**) muscle 6 (t(14)=18.60, p<0.0001, paired t test), (**G**) muscle 12 (t(14)=15.73, p<0.0001, paired t test) and (**H**) muscle 4 (t(15)=7.43, p<0.0001, paired t test). (I) Calculated EPSP ratios of 1b/1b+1s of m6, m12 and m4 (F(2,43)=26.03, p<0.0001, one-way ANOVA, Tukey post hoc test). Error bars indicate ±SEM. ****p<0.0001. n (NMJs/larva) is 15/12, 15/12, and 16/15 respectively. Calibration bar in **B** is 5μm.

In the larval neuromuscular circuit, 1b and 1s MNs are required for normal locomotion. Each MN type also has unique electrophysiological properties: 1b MNs display a higher basal neurotransmitter release probability (spontaneous activity, *P_r_*) but the quantal size is smaller compared to 1s MNs (Newman et al., 2017; Nguyen and Stewart, 2016). In traditional NMJ electrophysiology experiments, excitatory postsynaptic potentials (EPSPs) in muscles represent simultaneous stimulation of both neurons. Previous studies isolated 1b-derived and 1b+1s-derived EPSPs from different muscles (Table 1), and demonstrated that convergent 1b and 1s inputs do not equally contribute to the EPSPs recorded from different muscles (Aponte-Santiago et al., 2020; Genc and Davis, 2019; Kurdyak et al., 1994; Li et al., 2002; Lnenicka and Keshishian, 2000). Several forms of synaptic plasticity have been observed in the larval NMJ including facilitation and homeostatic compensation (Hallermann et al., 2010). As each muscle is co-innervated and individual synaptic activities can be distinguished, this system provides an ideal platform to investigate structural and physiological plasticity changes that enable one input to compensate for perturbations of a converging input on the same target.

**Table 1.** comparative 1b and 1s physiology from prior NMJ studies.

| muscle | 1b EPSP(mV) | 1s EPSP(mV) | 1b+1s EPSP(mV) | [Ca<sup>2+</sup>] | reference |
| --- | --- | --- | --- | --- | --- |
| 1 | 11.8 | - | 24.2 | 0.3 mM | Aponte-Santiago et al., 2020 |
| 6 | 14.9 | - | 24.3 | 1.0 mM | Li et al., 2002 |
| 6 | ~10 | - | ~35 | 1.8 mM | Kurdyak et al., 1994 |
| 6 | 15.6 | 25.7 | - | 0.5 mM | Genc et al., 2019* |
| 6 | 15.6 | - | 23.1 | 1.5 mM | Lnenicka and Keshishian, 2000 |
| 7 | 12.1 | - | 15.5 | 1.5 mM | Lnenicka and Keshishian, 2000 |

\* This study used optogenetics, instead of voltage injection, to differentially activate 1b or 1s.

In this study, we examined three separate muscles as each muscle is innervated by a unique combination of 1b and 1s inputs. First, we confirmed muscle-specific differences in 1b activities. Next, we removed 1s NMJs after innervation and examined 1b NMJ responses. All 1b inputs display structural and functional compensation after ablation of 1s NMJs, and some 1b inputs can partially compensate the loss of 1s evoked neurotransmission, confirming 1b synaptic plasticity in this system. In contrast, robust 1b synaptic plasticity does not occur if the 1s NMJ is never formed, suggesting that 1s co-innervation may establish an activity set point. Because most postsynaptic cells are multi-innervated, similar strategies may be utilized in other circuits.

## MATERIALS AND METHODS

### Fly strains

Fly strains:
| Fly strain | Source |
| --- | --- |
| *w<sup>1118</sup>* | (Carrillo et al., 2015) |
| *A8-GAL4* | (Venkatasubramanian et al., 2019) |
| *10XUAS-mCD8::GFP* | Bloomington Drosophila Stock center #32184 |
| *10XUAS-mCD8::GFP, A8-GAL4* | This paper |
| *DIP-α<sup>1-178</sup>* | (Xu et al., 2018) |
| *UAS-hid, rpr* | (Zhou et al., 1997) |
| *MHC-CD8::GCaMP6f-Sh* | (Newman et al., 2017) |

### Antibodies

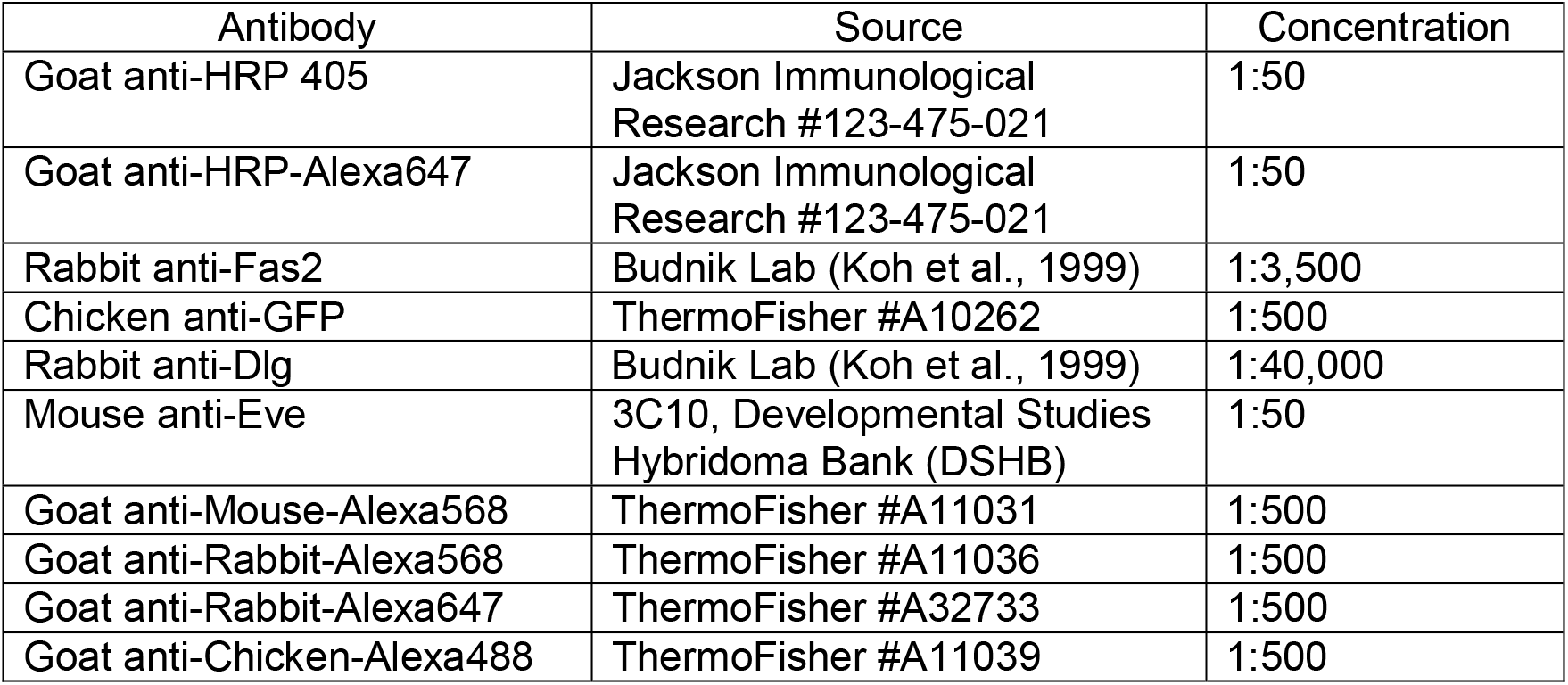

### Dissection and Immunocytochemistry

Embryonic dissections were performed as previous described (Lee et al., 2009). Briefly, egg laying chambers were setup with adult flies (15-20 females and 10-15 males) and grape juice plates (3% agar, 1.3% sucrose, 25% grape juice in water). After six-hour laying periods, grape plates covered in embryos were collected. Embryos were staged on double sided tape using the autofluorescence and shape of the gut (Bownes, 1975; Hartenstein et al., 1987) under a Zeiss V20 stereoscope, and then dechorionated with a sharpened metal probe and placed on grape juice agar. Embryos were transferred to double-sided tape on a Superfrost Plus slides (ThermoFisher #22-037-246) with the dissecting area outlined by a PAP pen (Research Products International, #195506), and then covered with 0.22μm filtered phosphate buffered saline (PBS) (0.01M Sodium Phosphate, 150mM Sodium Chloride). Embryos were opened with an electrolytically sharpened tungsten wire, removed from the vitellin membrane and then adhered to the charged slide. Dissected embryos were washed once with PBS, and then fixed for 30 minutes at room temperature using 4% paraformaldehyde (Electron Microscopy Sciences). Samples were then washed three times in 0.05% PBST (PBS with 0.05% TritonX100), and then blocked for 1 hour in 1% normal goat serum (1% Goat serum in 0.05% PBST). Samples were incubated in primary antibody solutions overnight at 4°C and washed three times in PBST. Samples were then incubated with secondary antibody solutions at room temperature for 2 hours and washed three times with PBST. Samples were finally mounted in vectashield (Vector Laboratories) and the coverslip sealed with nail polish.

First and third instar larval dissections were performed as previously described (Ashley et al., 2019). Wandering third instar larvae were dissected in PBS on Sylgard dishes and pinned down using sharpened 0.1mm Insect Pins (FST #26002-10). For first instar larvae, electrolytically sharpened tungsten pins were used to accommodate the size of smaller larvae. Samples were then fixed for 30 minutes using 4% paraformaldehyde and then transferred to 0.5ml Eppendorf tubes. Samples were blocked and treated with primary and secondary antibodies as embryo samples above. All larval washes and antibody incubations were performed with mild agitation on a nutator.

### Image Acquisition

All imaging was acquired on a Zeiss LSM800 confocal microscope with either a 20X plan-apo 0.8NA objective, a 40X plan-neofluar 1.3NA objective, or a 63X plan-apo 1.4NA objective. Laser intensity, pinhole and gain were adjusted to increase the signal but avoid overexposure. All samples from the same experiment were imaged under identical conditions. Example images are the maximum projection of the corresponding confocal Z stack (ImageJ).

### Image Analysis

#### dCE and vCE identification

Existence of vCE and dCE motor neurons were confirmed in embryo and first instar samples. dCE motor neurons were identified by the expression of GFP, Eve and their positions in the ventral nerve cord (VNC). vCE motor neurons were identified by the expression of GFP and their positions in the VNC. Final confirmation was done by identifying the muscle innervation patterns in larval abdominal hemisegments.

#### 1b bouton counting

1b bouton counting was performed in third instar samples. Boutons were examined using HRP and scored as 1s or 1b based on Dlg signal, as 1s boutons have a smaller and dimmer Dlg signal than 1b boutons. Satellite boutons were identified as small buds-like structures emerging from “parent” boutons (Lee and Wu, 2010).

### Electrophysiology and Analysis

All recordings were performed as previously described (Meng et al., 2020). Third instar larvae were dissected in modified HL3 saline (70 mM NaCl, 5 mM KCl, 10 mM MgCl2, 10 mM NaHCO3, 5 mM Trehelose, 115 mM Sucrose, 5 mM HEPES) with 0.3 mM calcium. Segmental nerves were cut near the ventral nerve cord to remove VNC input and then the larval fillet was perfused with modified HL3 saline containing 0.5 mM calcium. Body-wall muscle 6, 12 or 4 in abdominal segment 3 were impaled with 15-30 MΩ sharp electrode filled with 3 M KCl and recorded for miniature excitatory postsynaptic potentials (mEPSPs) for 2 minutes. Nerves were drawn into a suction electrode and stimulated to elicit excitatory postsynaptic potentials (EPSPs). Specifically, for muscle 6 and 12 EPSP recording, the whole segmental nerve bundle was stimulated whereas for muscle 4 EPSP recording, the branched nerve above muscle 5 were sucked to get a better stimulation. For each muscle, 24 EPSPs were recorded at 0.2 Hz and the largest 12 EPSPs were averaged to indicate the mean EPSP. Signals were amplified using a MultiClamp 700B Molecular Devices (MD) and digitized using a Digidata 1550B (MD). Stimulus was triggered via a Master‐9 stimulator (A.M.P.I.). Data was acquired using pClamp 10 software (MD) and analyzed using Mini Analysis software (Synaptosoft).

Analysis was only performed on muscle cells with resting potentials below −60mV. Quantal content was calculated by dividing the mean EPSP amplitude by the mean mEPSP amplitude for each muscle, and the resulting numbers pooled for each genotype. mEPSP amplitudes from the same genotype were pooled together and binned with 0.01 to calculate the cumulative probability or binned with 0.1 to show the event distribution.

### GCaMP Imaging coupled with Electrophysiology

Third instar larvae of *MHC-CD8::GCaMP6f-Sh* were dissected in modified HL3 saline with 0.5 mM calcium and followed the same procedure of electrophysiology. Larval fillets were visualized by Nikon FS microscope using a 40× long‐working distance objective and GCaMP positive 1b and 1s boutons of muscle 6, 12 and 4 were illuminated by an Aura II solid state illuminator. Together with electrophysiology stimulation, each stimulus triggered a GCaMP firing event with the corresponding EPSP. GCaMP signals were scanned and recorded by a PCO Edge 4.2 camera and NIS element software. Electrophysiology data was collected as described above. Given that 1s neurons have a lower evoked threshold than 1b neurons, stimulating voltage was tuned to isolate 1b alone firing events and 1b+1s firing events (Movie 1). If a stimulation only triggered GCaMP firing event at 1b boutons but not 1s boutons, the corresponding EPSP was counted as 1b alone EPSP. If a stimulation triggered GCaMP firing events at both 1b and 1s boutons, the corresponding EPSP was counted as 1b+1s EPSP. For each sample, 1b alone EPSPs and 1b+1s EPSPs were averaged respectively. 1b alone/1b+1s was calculated by dividing the mean 1b alone EPSP by the mean 1b+1s EPSP.

### Experimental design and statistical analysis

All statistical analysis was performed using Prism 8 software (Graphpad). Average and standard error of the mean (SEM) are reported. Outliers are determined by Q-test and excluded from the sample pools. For each data point at least eight animals per genotype were dissected and at least two biological replicates were examined. All data was assumed to follow a Gaussian distribution. Comparisons were performed by student t-test (Welch’s correction was used in cases of unequal variance), one-way ANOVA followed by Tukey test, or Kolmogorov-Smironv test (K-S test).

## RESULTS

### 1b and 1s inputs contribute to postsynaptic activity in a target-specific manner

Converging inputs contribute to the overall postsynaptic response. In this study, we asked to what extent one input can influence a convergent input’s structure and function. To address this, we first determined the activity contribution of each input on the postsynaptic muscle target in a wild-type condition. We chose muscles 6, 12 and 4 (m6, m12 and m4) because 1) prior studies showed that each 1b contributes a unique percentage of the total postsynaptic activity, 2) these muscles have been frequently analyzed in NMJ studies (Menon et al., 2013; Nose, 2012), and 3) the 1b and 1s innervation patterns on these muscles (all muscles have unique 1b MNs but m6 and m12 are innervated by the same 1s MN) enabled *identification* of common and muscle-specific principles.

We measured the 1b MN contribution to the total EPSP on a muscle by muscle basis. 1b and 1s axons fasciculate into nerve bundles as they exit the ventral nerve cord (VNC), which impeded us from physically stimulating each neuron independently without patching each cell body (Choi et al., 2004). To circumvent this, we combined NMJ electrophysiology with a postsynaptically-targeted genetically encoded calcium indicator, SynapGCaMP6f (*MHC-CD8-GCaMP6f;* (Newman et al., 2017)). Some of the earliest evidence for the existence of two distinct MN populations was uncovered over 40 years ago (Jan and Jan, 1976) through observations that different voltage injections elicited two populations of muscle EPSPs. It was later found that lower voltage injection generated action potentials in 1b MNs (Lnenicka and Keshishian, 2000), therefore we varied the stimulus protocol to independently elicit and record 1b-and 1b+1s-derived EPSPs (see Materials and Methods). SynapGCaMP6f fluorescence changes at 1b and 1s NMJs confirmed whether the recorded EPSPs were due to 1b alone or 1b+1s activity (Figure 1B, Movie 1). Using this procedure, the average combined EPSP amplitude (1b+1s) in m6 was 31.55 mV and the average 1b-derived EPSP was 17.17mV (Figure 1C,F). Thus, the m6-1b input contributes 54% of the combined m6 EPSP (Figure 1I). Interestingly, at m12 the 1b input contributed 31% of the combined EPSP, and at m4, the 1b represents 62% (Figure 1D,E,G-I). Thus, we determined the contribution of each input (1b and 1s) to the postsynaptic muscle activity in wild type larvae and found that the relative strength of each input differed between muscles, with the m4-1b MN contributing the most and the m12-1b the least. Notably, m6-1b and m12-1b contributions are different even though they are innervated by the same 1s MN. These data establish a model in which to introduce perturbations and examine synaptic plasticity.

### Cell-specific genetic ablation of 1s MNs by ectopic *hid,rpr* expression

To begin to examine if one MN can respond to perturbations in a convergent MN, we needed to disrupt one input and monitor the impact on the converging input. Additionally, being able to disrupt one input before or after innervation can shed light on whether co-innervation is required for synaptic plasticity. We needed to identify drivers that explicitly express within subsets of the converging neurons. In a previous study, we showed that *DIP-α* is expressed in subsets of neurons in the larval VNC, including many interneurons and two 1s MNs, and that removal of *DIP-α* impeded 1s innervation of m4 (Ashley et al., 2019). To gain genetic access specifically to the 1s MNs, we examined a GAL4 driver derived from the *DIP-α* promoter. In the adult neuromuscular circuit, this driver (hereafter referred to as *A8-GAL4*, or A8) was found to express in a small subset of MNs (Venkatasubramanian et al., 2019). We used *A8-GAL4* to drive UAS-GFP and found expression in two pairs of segmentally repeated neurons in the third instar VNC (Figure 2A and 2B, arrows). The labeled neurons located ventrally in the VNC have axons that project medially and dorsally towards the neuropil (Figure 2-1A, arrow). The other A8-expressing neurons are located in the dorsal region of the VNC and showed an ipsilateral projection with a large dendritic arbor (Figure 2-1B, arrowhead) and an axon that exits the VNC (Figure 2B and Figure 2-1B, asterisks). The dorsal cells were co-labeled with Even-skipped (Eve), which labels three medial neurons, aCC, pCC, and MNISN-1s (Broadus et al., 1995; Doe et al., 1988). Based on the location of these *A8* positive cells, these are the MNISN-1s neurons (also called the dorsal common exciter, dCE) (Figure 2B). Examination of nerves exiting the VNC shows two axons in each nerve (Figure 2B and Figure 2-1B, asterisks), suggesting that these *A8* positive neurons are motor neurons. *Drosophila* larval MNs make connections with their muscle targets with very high fidelity, allowing us to unequivocally determine the identity of the MNs based on their innervation pattern. In A8>GFP third instar larvae, one axon innervates the ventral muscles including muscles 6, 7, 12, and 13, similar to the connectivity pattern of MNISNb/d-1s (also called the ventral common exciter, vCE; Meng et al., 2020) (Figure 2C). The other axon innervates the dorsal muscles, corresponding to dCE (Figures 1A and 2D). In order to distinguish 1b and 1s NMJs, we also stained for Discs large (Dlg) as 1b NMJs have significantly more DLG (Guan et al., 1996) (Figure 2C and 2D). *A8* is not expressed in the third 1s MN that innervates lateral transverse muscles. Importantly, *A8* does not label any other MNs but only a few additional cells in one segment of the VNC (Figure 2A, carets). In summary, *A8* labels two 1s MNs (the vCE and dCE) within each hemisegment.

**Figure 2.**
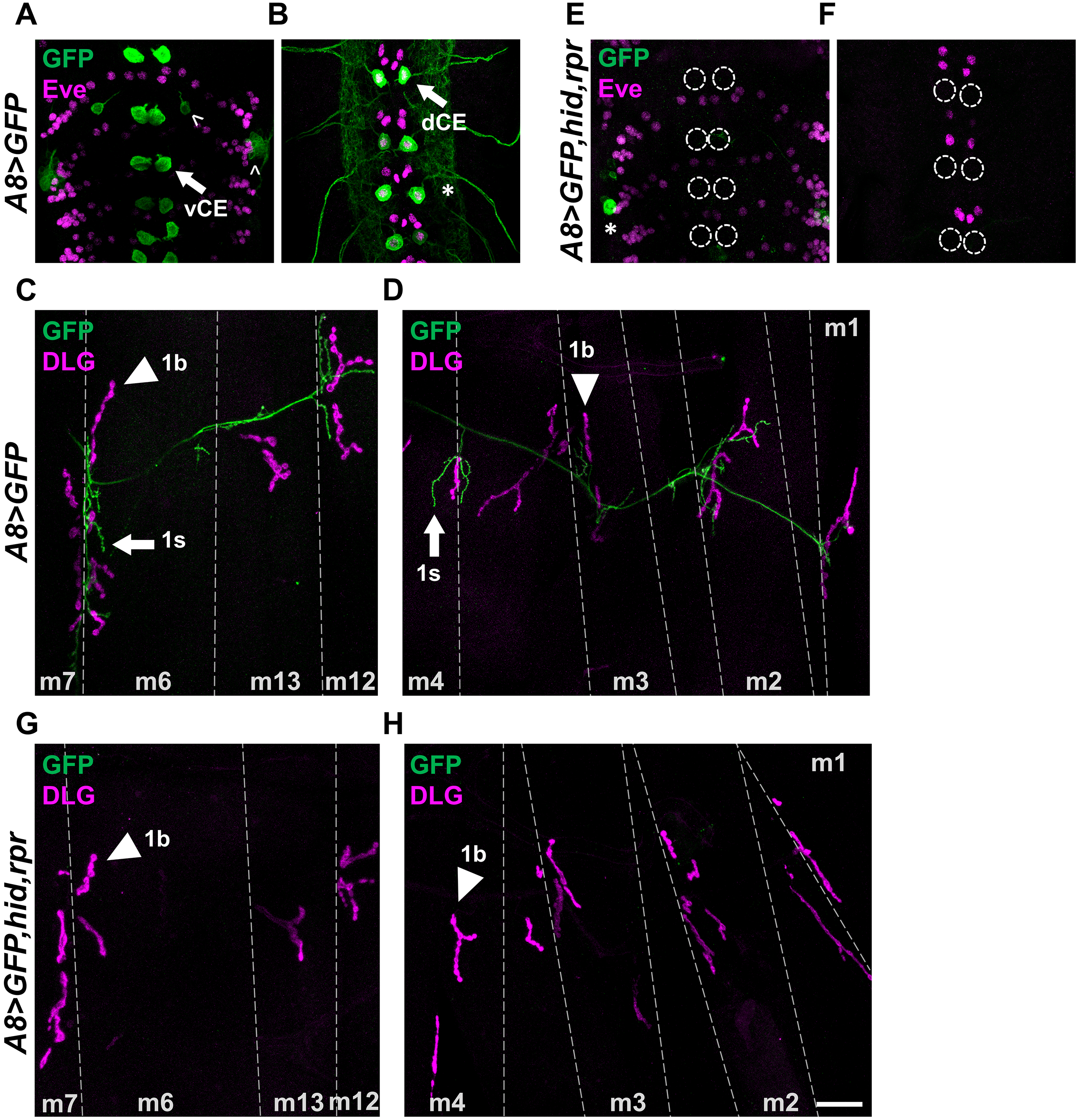
A8-GAL4 drives expression in 1s MNs and can be used to ablate 1s MNs. (**A**, **B**) Representative 3rd instar larval ventral nerve cord Z-sections showing (**A**) ventral and (**B**) dorsal cell bodies labeled with GFP (green) and Eve (magenta, labels nuclei of dCE and other neurons), in A8>GFP. Arrows indicate GFP positive vCE and dCE cell bodies. Carets indicate other cells that express A8. Asterisks indicate two axons exiting the VNC (See also Figure 2-1). (**C**, **D**) Representative 3rd instar larval abdominal hemisegment labeled with GFP (green) and the postsynaptic marker DLG (magenta), in A8>GFP. (**C**) Ventral muscles (m6, m7, m13 and m12) innervated by vCE and (**D**) dorsal muscles (m4, m3, m2 and m1) innervated by dCE. Arrows indicate 1s NMJs and arrowheads indicate 1b NMJs. (**E**, **F**) Representative 3rd instar larval VNCs lacking (**E**) vCE and (**F**) dCE labeled with GFP (green) and Eve (magenta) in A8>GFP,hid,rpr. Dashed circles mark the absence of vCE and dCE. Note that both vCE and dCE cell bodies are ablated by the third instar stage. Asterisk marks a GFP positive cell that remained. (**G**, **H**) Representative 3rd instar larval abdominal hemisegment showing (**G**) ventral muscles and (**H**) dorsal muscles, labeled with GFP (green) and DLG (magenta) in A8>GFP,hid,rpr. Note all NMJs from vCE and dCE are absent (no GFP). Arrowheads indicate 1b NMJs. Calibration bar in **A,B,E,F** is 27μm and **C,D,G,H** is 35μm.

We used *A8-GAL4* to drive *head involution defective (hid)* and reaper (*rpr*) in vCE and dCE. *hid* and *rpr* have important functions in programmed cell death, and ectopic expression of both genes more reliably induces neuronal death than either gene alone (Pauls et al., 2015; Zhou et al., 1997). As shown in Figure 2C and 2D, A8>GFP robustly labels 1s NMJs but in third instar larvae that ectopically express *hid,rpr* (*A8>GFP,hid,rpr*), all vCE or dCE cell bodies and NMJs are absent (Figure 2E-H). Thus, expression of *hid,rpr* is sufficient to genetically ablate 1s MNs.

To determine whether cell death occurred prior to or after innervation, we examined earlier developmental stages and visualized GFP in 1s MN cell bodies and NMJs in A8>GFP and *A8>GFP,hid,rpr*. Neuromuscular innervation is established at embryonic stage 16 (Broadie and Bate, 1993; Halpern et al., 1991; Yoshihara et al., 1997), so we focused on stage 15 and later stages including stage 17 and first instar larvae. In stage 15 embryo controls (A8>GFP), only a subset of 1s MNs are detected since not all dCE cell bodies (Eve positive) are co-stained with GFP and no vCE cell bodies are observed (Figure 3A and 3B). Age matched *A8>GFP,hid,rpr* embryos showed similar GFP expression patterns to controls (Figure 3C and 3D); thus, no cell death occurred before neuromuscular innervation. By embryonic stage 17, more vCE and dCE cell bodies expressed *A8* but the lack of GFP in some cells (Figure 3E and 3F) suggests that *A8* expression has not reached maximal levels. In stage 17 *A8>GFP,hid,rpr* embryos, cells are undergoing apoptosis as revealed by significant decreases in GFP and by loss of Eve staining (Figure 3G and 3H); Eve is a transcription factor found in the nucleus and its loss suggests nuclear degradation. Finally, in A8>GFP first instar larvae, all vCE and dCE cell bodies were labeled (Figure 3I and 3J)) and NMJs were present (Figure 3M). Age-matched *A8>GFP,hid,rpr* larvae completely lacked vCE and dCE cell bodies (Figure 3K and 3L), although some 1s NMJs were observed (Figure 3N), suggesting that 1s NMJ are established before ablation in this genetic background. Given that neuromuscular innervation happens at embryonic stage 16 (Broadie and Bate, 1993; Halpern et al., 1991; Yoshihara et al., 1997), *A8>GFP,hid,rpr*-induced genetic ablation provides a means to specifically target 1s MNs for cell death after establishing NMJs. Thus, *A8-GAL4* is specifically expressed in vCE and dCE motor neurons and driving ectopic expression of cell death genes with *A8* triggers apoptosis in 1s MNs after synaptic contacts are made. Therefore, *A8>GFP,hid,rpr*-induced genetic ablation provides a tool to eliminate 1s MNs after NMJs are established.

**Figure 3.**
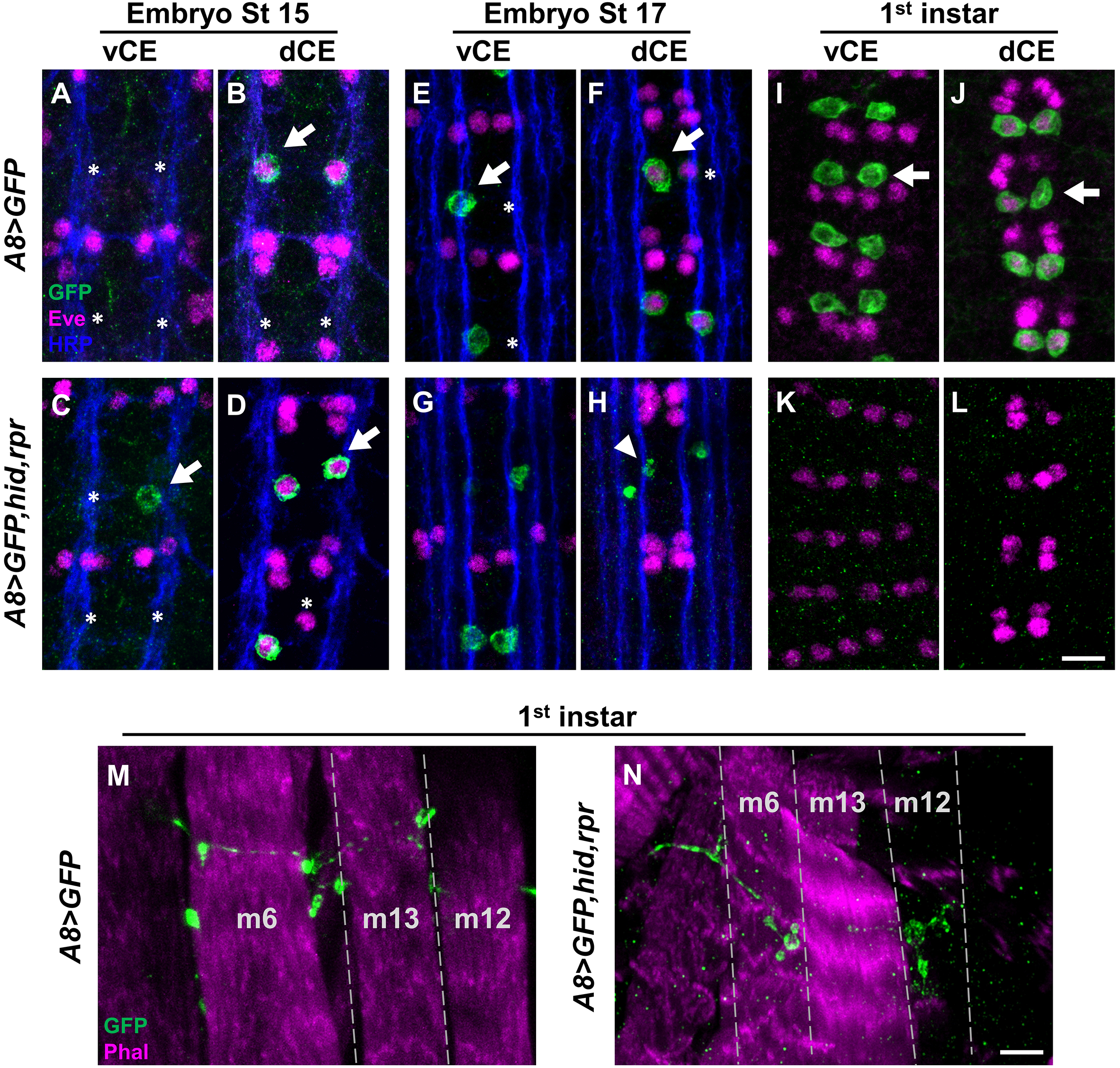
A8>hid,rpr-induced cell death occurs after 1s innervation. (**A-L**) Representative images depicting the presence or absence of vCE and dCE cell bodies from (**A-D**) embryonic stage 15, (**E-H**) stage 17 and (**I**-**L**) first instar larval VNCs labeled with GFP (green), Eve (magenta) and Fasciclin 2 (blue) in control (A8>GFP) and mutant (A8>GFP,hid,rpr) animals. Arrows and asterisks indicate the cells expressing or not expressing A8, respectively. A8 expression begins at (**A**,**B**) embryonic stage 15. In A8>GFP,hid,rpr, vCEs and dCEs undergo apoptosis starting at (**G**,**H**) embryonic stage 17, noted by the loss of Eve staining in dCE and GFP positive debris (indicated by arrowhead), and are completely absent in (**K**,**L**) first instar larvae. (**M**,**N**) Representative 1s NMJs labeled with GFP (green) and a muscle marker, phalloidin (magenta), in (**M**) control and (**N**) mutant first instar larvae. Note 1s NMJs are labeled by GFP in control animals and some 1s boutons are still present in mutants, suggesting A8>GFP,hid,rpr-induced cell death happens after 1s NMJ formation. Calibration bar in **A**-**F** is 9μm, and **G**,**H** is 4μm.

### The 1b NMJ expands upon loss of 1s synapses

NMJ structural plasticity can be induced by perturbation of synaptic function (Budnik et al., 1990; Goel et al., 2019; Jarecki and Keshishian, 1995; Perry et al., 2020; Sigrist et al., 2003). Here, we examined whether loss of a synaptic input can modulate structural changes at an adjacent synaptic arbor. The size of each of the examined arbors is well characterized (represented by the number of boutons) and this allows us to observe structural changes due to perturbations. We genetically ablated 1s MNs (vCE and dCE) and counted the number of boutons to estimate the size of 1b arbors on m6, m12, and m4 in wandering third instar larvae (NMJ expansion is complete). Loss of 1s innervation on all three muscles revealed an increase in the number of 1b boutons (Figure 4A-G), suggesting that 1b NMJs expand when adjacent 1s synapses are eliminated. Interestingly, we also observed an increase in small budding boutons, called satellite boutons, emanating from 1b boutons (Figure 4D-F, insets, and 4H). These structures represent immature but functional boutons as they contain all synaptic machinery and postsynaptic receptors (Carrillo et al., 2015; Dickman et al., 2006; Lee and Wu, 2010; O’Connor-Giles and Ganetzky, 2008; O’Connor-Giles et al., 2008; Torroja et al., 1999). Thus, all 1b MNs initiate structural plasticity mechanisms to respond to loss of adjacent 1s inputs.

**Figure 4.**
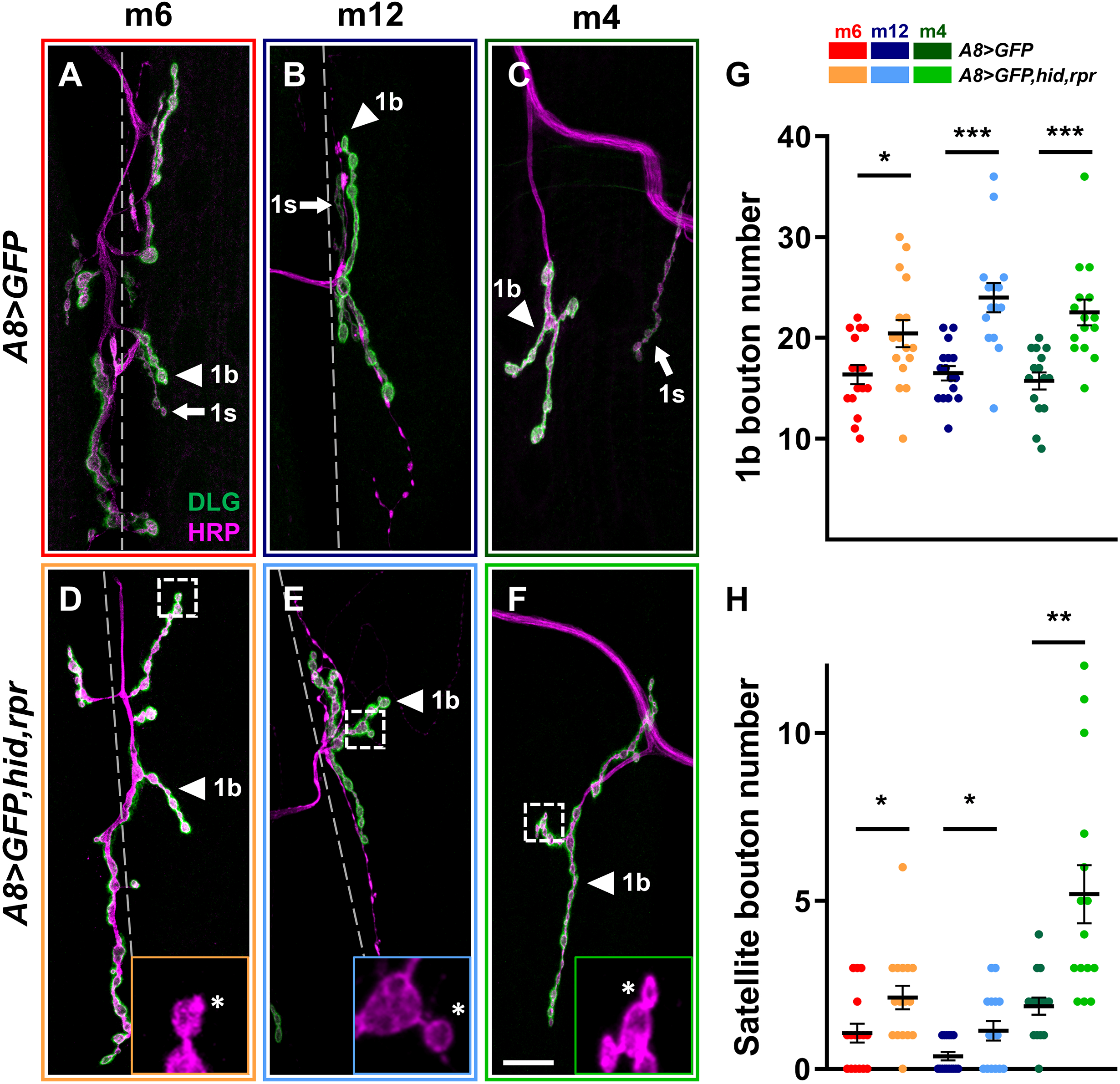
1b synaptic arbors expand upon on loss of 1s MNs. (**A-F**) Representative NMJ arbors (1b arbors and 1s arbors, arrowheads and arrows, respectively) from (**A**, **D**) m6, (**B**, **E**) m12 and (**C**, **F**) m4 labeled with Dlg (green) and HRP (magenta) in control (A8>GFP) and mutant (A8>GFP,hid,rpr) third instar larvae, respectively. (**D**-**F**) Insets show satellite boutons, indicated by asterisks. Note 1s NMJs are absent in mutant animals. (**G**) Quantification of 1b bouton number of m6 (t(30)=2.458, p=0.02, unpaired t test), m12 (t(26)=4.449, p=0.0001, unpaired t test) and m4 (t(28)=4.431, p=0.0001, unpaired t test) in control and mutant animals (satellite 1b boutons were not included). (**H**) Quantification of satellite boutons from m6 (t(30)=2.359, p=0.025, unpaired t test), m12 (t(26)=2.397, p=0.0269, unpaired t test with Welch’s correction) and m4 (t(28)=3.682, p=0.0019, unpaired t test with Welch’s correction) in control and mutant animals. Error bars indicate ±SEM.

### Loss of 1s innervation leads to 1b synaptic plasticity

To determine to what extent a synaptic input can influence a converging input’s functional synaptic plasticity, we examined 1b spontaneous and evoked activities at muscles where 1s MNs are ablated after innervation. 1b and 1s MNs have unique basal neurotransmitter release properties. For example, 1b-derived spontaneous events (stimulus-independent basal release of neurotransmitter vesicles; also referred to as mini EPSPs, mEPSPs), have smaller amplitudes compared to 1s mEPSPs (Newman et al., 2017; Nguyen and Stewart, 2016). Therefore, ablation of 1s inputs should shift the mean mEPSP amplitude towards the smaller 1b-like amplitude if there is no compensation. We performed current clamp recordings from m6, m12 and m4. Indeed, *A8>GFP,hid,rpr* revealed decreased mEPSP amplitudes compared to A8>GFP controls, and a significant shift in cumulative amplitude probability distribution (Figure 5A-G). Due to the inability to assign spontaneous events as either 1b or 1s in controls, we are unable to unambiguously determine if the 1b mEPSP amplitudes are affected by the loss of 1s inputs. Nonetheless, we can conclude 1b MNs cannot fully restore the average mEPSP amplitudes to wild type levels.

**Figure 5.**
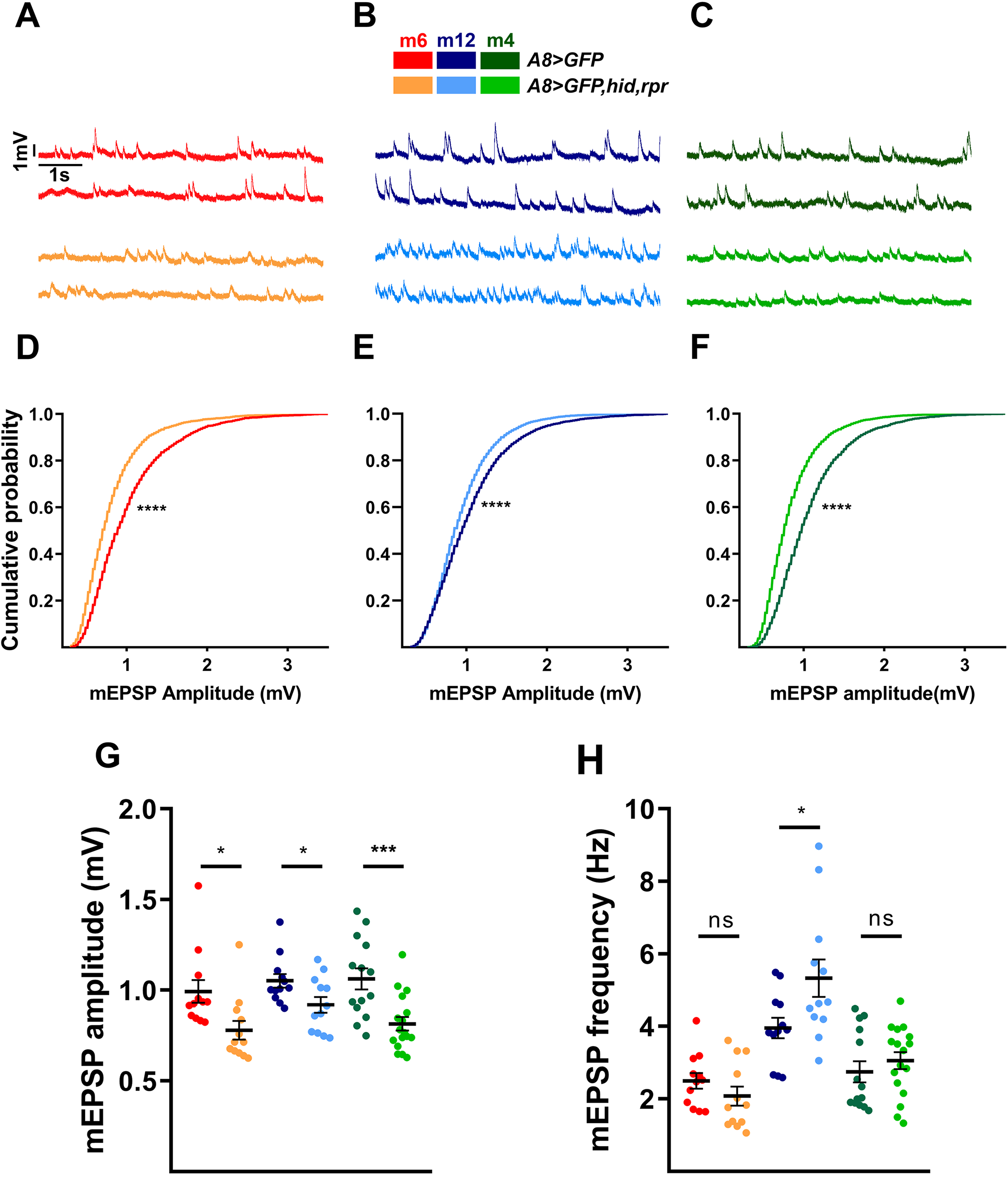
Loss of 1s MNs decreases overall mEPSP amplitudes and increases 1b mEPSP frequencies. (**A**-**C**) Representative mEPSP recordings from (**A**) m6, (**B**) m12 and (**C**) m4 in control (A8>GFP) and mutant (A8>GFP,hid,rpr) animals. Traces are color-coded as indicated in the color key (**G**). (**D-F**) Pooled cumulative probability distributions from (**D**) m6 (p<0.0001, K-S test), (**E**) m12 (p<0.0001, K-S test), (**F**) m4 (p<0.0001, K-S test). (**G**) Quantification of mEPSP amplitude of m6 (t(22)=2.630, p=0.0153, unpaired t test), m12 (t(22)=2.294, p=0.0317, unpaired t test) and m4 (t(29)=3.700, p=0.0009, unpaired t test) in controls and mutants. Each data point represents the average mEPSP amplitude from one sample. (**H**) Quantification of mEPSP frequencies of m6 (t(22)=1.224, p=0.2339, unpaired t test), m12 (t(22)=2.331, p=0.0293, unpaired t test) and m4 (t(29)=0.8369, p=0.4095, unpaired t test) in controls and mutants. Each data point represents the average mEPSP frequency from one sample. Error bars indicate ±SEM. *p<0.05, **p<0.01, ***p<0.001, ****p<0.0001. n (NMJs/larva) is 12/9, 12/11, 12/9, 12/12, 14/11, and 17/14 respectively for **D-H**.

Another measure of basal activity is the rate of spontaneous neurotransmitter release. In prior studies, mEPSP frequencies were found to be higher at 1b synapses than 1s synapses (e.g., 2.3 Hz at m4-1b synapses and 1 Hz at m4-1s synapses; Newman et al., 2017); thus, if elimination of 1s inputs does not affect 1b baseline activity, overall mEPSP frequencies should decrease about one third. However, we did not observe any reduction of mEPSP frequencies on m6, m12, and m4 when comparing A8>GFP and *A8>GFP,hid,rpr* animals, and m12 even showed an increased rate of spontaneous activity (Figure 5A-C and 5H). Importantly, these spontaneous events correspond to 1b-derived mEPSPs since a) the mean amplitudes decreases (Figure 5G), and b) there are no 1s boutons observed at this stage (Figure 2G and 2H). These results strongly suggest that 1b MN synaptic plasticity mechanisms MN detect the loss of co-innervating 1s MNs and elevate their spontaneous firing rate.

Having demonstrated 1b synaptic plasticity of basal activity, we next examined whether 1b stimulus-dependent activity could also be modified by the loss of 1s innervation. Unlike spontaneous neurotransmitter release, EPSPs require stimulation to depolarize the presynaptic neuron above threshold. This suprathreshold stimulation triggers an action potential (AP) to induce neurotransmitter release and elicit a postsynaptic response. In order to examine if 1b inputs can compensate for the loss of 1s synaptic drive, we recorded EPSPs in both A8>GFP and *A8>GFP,hid,rpr* (Figure 6A-F). To determine if 1b-derived EPSP amplitudes are affected by loss of 1s inputs, we normalized the EPSPs (*A8>GFP,hid,rpr* EPSP/A8>GFP EPSP) and compared these values to the calculated 1b/1b+1s ratio in Figure 1I. This analysis allows for the direct comparison of 1b-derived EPSPs with and without convergent 1s inputs.

**Figure 6.**
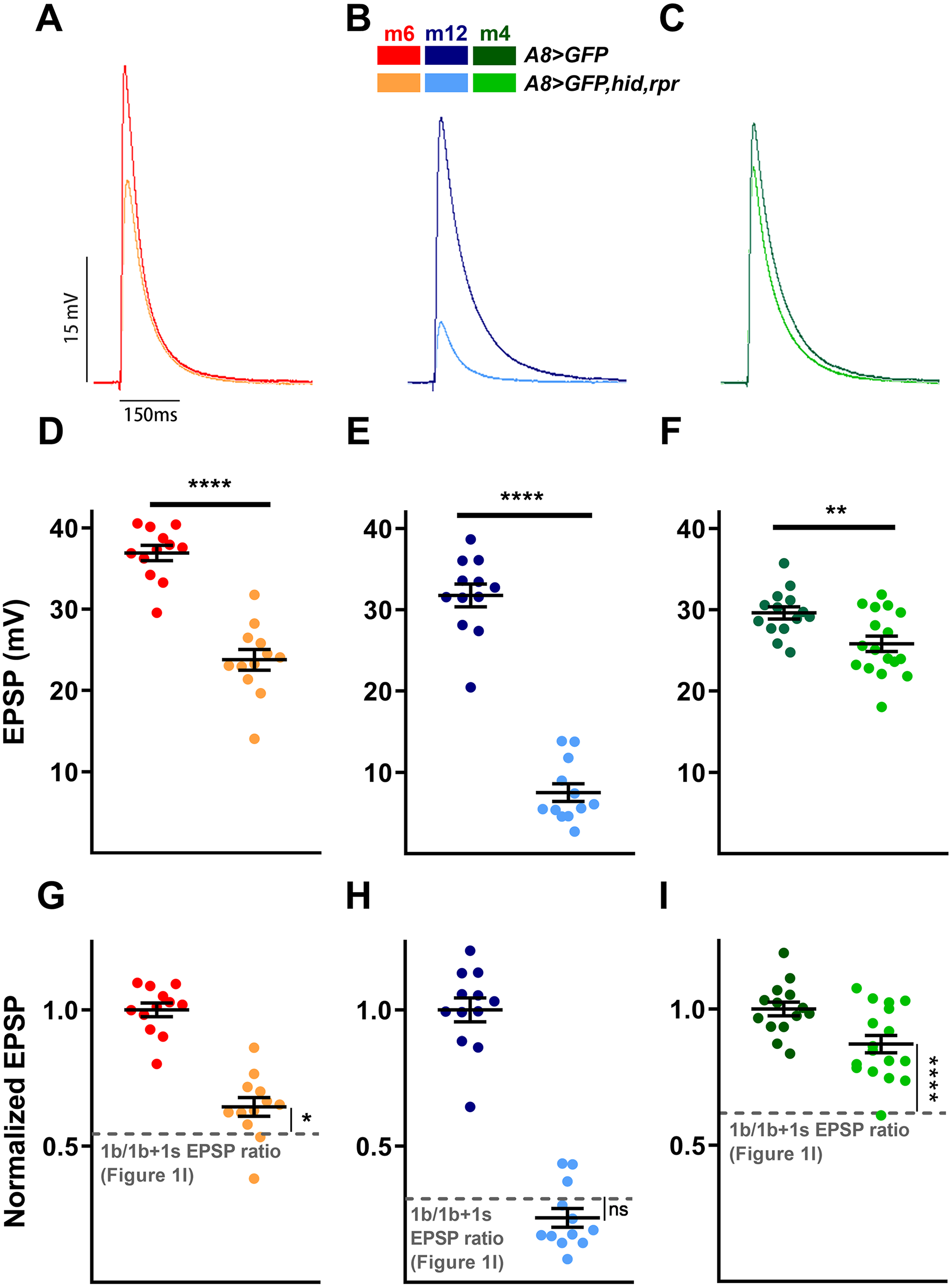
Some 1b MNs elevate evoked EPSPs in a target-specific manner in the absence 1s inputs. (**A**-C) Representative EPSP traces of 1b+1s and 1b alone on (**A**) m6, (**B**) m12, and (**C**) m4. (**D**-**F**) Quantification of EPSP amplitudes in (**D**) m6 (t(22)=8.306, p<0.0001, unpaired t test), (**E**) m12 (t(22)=13.82, p<0.0001, unpaired t test) and (**F**) m4 (t(29)=3.057, p=0.0048, unpaired t test) in controls (A8>GFP) and mutants (A8>GFP,hid,rpr) (See Figure 6-1 for quantal content). (**G**-**I**) Normalized EPSPs from (**G**) m6 (t(25)=2.301, p=0.0300, unpaired t test), (**H**) m12 (t(25)=1.552, p=0.1332, unpaired t test) and (**I**) m4 (t(31)=4.605, p<0.0001, unpaired t test) in control and mutant third instar larvae. Average combined EPSP (1b+1s) from control is used for normalization. Normalized mutant EPSPs are compared to the EPSP ratio of 1b/1b+1s calculated from corresponding muscles in Figure 1I, indicated by grey dashed line. Note m6-1b slightly compensates, m12-1b does not compensate and m4 largely compensates the loss of 1s-derived EPSPs. Error bars indicate ±SEM.

We observed muscle-specific changes in 1b-derived EPSPs (Figure 6G-I). At m4, we observed a significant increase in 1b-derived EPSPs compared to the control 1b/1b+1s ratio (Figure 6I). By repeating the analysis at other muscles, we found a mild increase in 1b AP-induced EPSPs at m6 (Figure 6G), and surprisingly, no change at m12 (Figure 6H). When comparing these EPSP compensations (Figure 6G-I) to the wild type 1b contributions at each muscle (Figure 1I), a pattern arises whereby the m4-1b has both the highest percentage of compensation and the largest 1b contribution. Meanwhile, m12-1b did not compensate and displayed the smallest 1b contribution.

Finally, the quantal content, a calculation of the approximate amount of neurotransmitter released per stimulation, at m4 was restored to control levels, whereas m6 and m12 quantal content was reduced (Figure 6-1A-C). Overall, when ablating 1s MNs after NMJ formation, 1b MNs that innervate the same postsynaptic muscle upregulated their rate of spontaneous release, and importantly, m6-1b and m4-1b MNs partially compensated the respective EPSPs to maintain near combined (1b+1s) levels.

### Robust 1b synaptic plasticity requires 1s co-innervation

Above, we examined the structural and functional synaptic plasticity of one neuron when a converging neuron is ablated after innervation (*A8>GFP,hid,rpr*). To probe if initial co-innervation is required for plasticity, we explored another context when the 1s MN never makes contact with the postsynaptic target. *DIP-α* is required for recognition of m4 by the dCE; loss of *DIP-α* results in the lack of 1s innervation on m4 (Ashley et al., 2019). We combined the *DIP-α^null^* mutant with the A8>GFP to monitor 1s innervation of m4. Loss of 1s NMJs was confirmed with Dlg staining (Figure 7A and 7B). Quantification of m4-1b boutons and satellite boutons in *DIP-α^null^* mutants showed no difference from controls (Figure 7C and 7D). Also, in *DIP-α^null^* mutants, m4 mEPSP frequency was significantly reduced (Figure 7E and 7F), in contrast to the 1s ablation (Figure 5H). These results suggest that the m4-1b NMJ failed to respond to the loss of 1s synapses in *DIP-α^null^* mutants. Next, we examined EPSPs (Figure 7G and 7H). Surprisingly, loss of 1s innervation in *DIP-α^null^* mutant larvae revealed only a slight compensatory increase in the 1b-derived EPSP amplitude on m4 (Figure 7I). Comparing *A8>GFP,hid,rpr* and *DIP-α^null^* mutants to their respective controls, we observed differential upregulation of normalized 1b activity: 45% and 23%, respectively. Finally, quantal content showed no compensation in *DIP-α^null^* mutants (Figure 7-1). Thus m4-1b synaptic plasticity is not as robust in *DIP-α^null^* larvae, compared to *A8>GFP,hid,rpr*. These data suggest that co-innervation by 1s serves an important role to establish an activity set point (see Discussion for our model).

**Figure 7.**
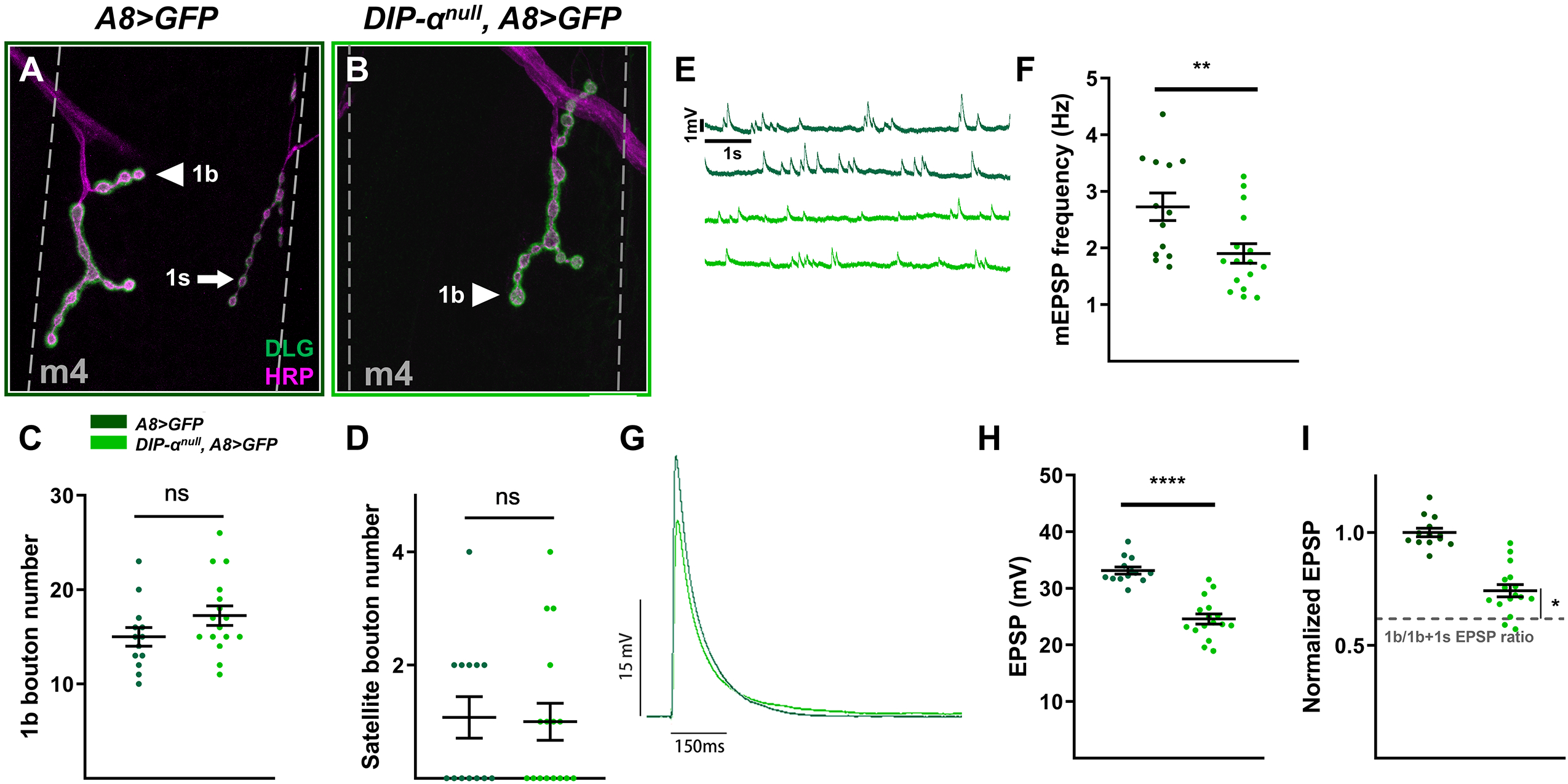
Robust 1b MN compensation requires co-innervation by a 1s MN. (**A,B**) Representative m4 NMJs labeled with DLG (green) and HRP (magenta) from (**A**) control (A8>GFP: dark green) and (**B**) mutant (DIP-αnull, A8>GFP: light green) animals. (**C**) Quantification of 1b boutons on m4 in control and mutant animals(t(27)=1.552, p=0.1323, unpaired t test). (**D**) Quantification of satellite boutons on m4 in control and mutant animals (t(27)=0.1563, p=0.8770, unpaired t test). (**E**) Representative mEPSP recordings of control and mutant larvae. (**F**) Quantification of mEPSP frequencies from m4 in control and mutant animals (t(27)=2.824, p=0.0088, unpaired t test). (**G**) Representative EPSP traces from control and mutant animals. (**H**) Quantification of EPSP amplitudes in control and mutant animals (t(27)=7.505, p<0.0001, unpaired t test) (See also Figure 7-1 for quantal content). (I) Normalized EPSPs from m4 in control and mutant animals (t(24.28)=2.358, p=0.0268, unpaired t test with Welch’s correction). Average combined EPSP (1b+1s) in control is used for normalization. Normalized mutant EPSP is compared to the EPSP ratio of 1b/1b+1s calculated from corresponding muscle in Figure 1I, indicated by grey dashed line. Note m4-1b-derived EPSP increases 23% in DIP-α mutants but increases 45% in A8>GFP,hid,rpr (Figure 6I). Error bars indicate ±SEM. *p<0.05, **p<0.01. n (NMJs/larva) is 13/8,16/8 for C,D and 13/10, 16/10 for E-I respectively. Calibration bar in A,B is 14μm.

## DISCUSSION

The major gap this paper addresses is to what extent one synaptic input can influence a converging input’s structural and functional plasticity. In this paper, we examine the *Drosophila* neuromuscular circuit and demonstrate 1b synaptic plasticity upon loss of converging 1s inputs. The muscles examined in this study, m6, m12, and m4, are co-innervated by 1b and 1s MNs. First, we established an activity baseline in wild type and uncovered that 1b MNs contribute a unique percentage of the total EPSPs in a muscle-specific manner. Genetic ablation of 1s MNs (vCE and dCE) after innervation led to expansion of 1b synaptic arbors and elevation of their basal spontaneous release. Furthermore, 1b MNs elevated evoked release in a target-specific manner. Therefore, we conclude that 1b MNs can structurally and functionally compensate the loss of 1s MNs. In larvae where the m4-1s NMJs never form, we observed no changes in m4-1b NMJ size or spontaneous release, and decreased EPSP compensation. These results suggest that co-innervation by 1b and 1s MNs may be required to establish an activity set point for robust compensation by 1b MNs in the absence of convergent 1s inputs. Thus, unidentified signaling pathways allow for inputs to respond to changes in adjacent synapses.

### Target-specific 1b MN synaptic weight and plasticity

Synaptic activity must be tightly regulated to extract meaningful information and produce an appropriate response. Homeostatic mechanisms constrain synaptic strength within an optimal range to combat perturbations in synaptic activity. In circuits such as the *Drosophila* larval NMJ, where each postsynaptic cell receives convergent inputs, it is not clear if every input contributes to these compensatory changes. Furthermore, can an input respond to perturbations caused by changes to an adjacent input? And if so, is this a general feature of all of presynaptic inputs and target cells?

In complex neural circuits, dissecting contributions of individual presynaptic inputs to the total postsynaptic activity, also referred to as the synaptic weight (Bhalla, 2008; Magee and Cook, 2000), remains difficult due to the thousands of inputs that can converge on a single cell. The relatively simple larval NMJ allows partitioning of synaptic inputs since each muscle is innervated by few MNs. Combining electrophysiology and calcium imaging, we found that 1b synaptic weights differ on m6, m12, and m4; thus confirming previous studies that examined m1, m6, and m7 (Aponte-Santiago et al., 2020; Genc and Davis, 2019; Kurdyak et al., 1994; Li et al., 2002; Lnenicka and Keshishian, 2000). This system served as ideal model to investigate how synaptic weights change in response to loss of adjacent inputs.

Examination of the 1b synaptic weights in animals with ablated 1s MNs revealed a muscle-specific pattern of 1b plasticity. Comparing this pattern to control 1b synaptic weights revealed that the degree of EPSP compensation upon removal of 1s inputs correlated with the degree of control 1b synaptic weights. Thus, robust 1b MNs that carry more of the synaptic drive may be endowed with certain synaptic plasticity mechanisms that respond to loss of adjacent inputs.

### A model for 1b MN synaptic plasticity

During larval development, muscle surface area increases ~100-fold, and to maintain synaptic efficacy due to the decrease in input resistance, the NMJ expands rapidly (Li et al., 2002; Menon et al., 2013; Ruiz-Canada and Budnik, 2006; Van Vactor and Sigrist, 2017; Zito et al., 1999). Remarkably, at the first instar stage, the EPSP maximum amplitude is reached and maintained through 4 days of larval development (Davis and Goodman, 1998; Li et al., 2002). One intriguing hypothesis is that 1b+1s co-innervation determines the EPSP set point during embryogenesis and this set point is maintained throughout development.

Conceptual and computational models of synaptic homeostasis rely on an activity set point to stabilize neurons when confronted with perturbations (Davis, 2013; LeMasson et al., 1993; Liu et al., 1998; O’Leary et al., 2014; Turrigiano, 2007). Each neuron must account for all presynaptic inputs to produce a defined output (i.e. the set point). The structural and functional properties of each input thus determine not only its contribution to the postsynaptic activity but also its ability to respond to perturbations in synaptic function. For example, transcription factors not only regulate the temporal expression of ion channels that shape neuronal excitability, but also homeostatic mechanisms (Davis, 2013; Diering et al., 2017; Engelmann and Haenold, 2016; Parrish et al., 2014; Schaukowitch et al., 2017; Turrigiano, 2007). Like many activity-dependent processes (Ataman et al., 2008; Berke et al., 2013; Carrillo et al., 2010; Vonhoff and Keshishian, 2017a), the optimal set point may be established during a narrow time-window of development. This hypothesis was tested in a *Drosophila* seizure mutant by inhibiting activity during embryonic development and observing suppression of seizures in postembryonic stages (Giachello and Baines, 2015). Thus, manipulating activity during an embryonic critical period may alter the activity set point.

We propose a model to describe how 1b MNs increase their arbor sizes and basal and evoked activities due to the loss of convergent 1s synapses. Functional NMJs begin to form during late embryonic development (Prokop et al., 1996; Vonhoff and Keshishian, 2017b; Yoshihara et al., 1997), and before the larva hatches, the neuromuscular innervation map is established. Since EPSP amplitudes are maintained throughout larval development, the set point is likely determined in late embryo/early first instar. Blocking the initial formation of 1s NMJs would create a set point that is devoid of 1s influence, and thus, the corresponding 1b input would reference this alternate set point. If the 1s MN is removed after synaptogenesis, the correct set point is established and the 1b responds accordingly.

### Potential mechanisms to regulate co-innervation plasticity

Homeostatic regulation can manifest in pre- and/or postsynaptic changes, including neurotransmitter release (Davis, 2013; Davis and Goodman, 1998; Davis and Muller, 2015; Frank et al., 2020; Petersen et al., 1997) and neurotransmitter receptor dynamics and abundance (Frank et al., 2020; Turrigiano et al., 1998; Wierenga et al., 2005), respectively. In our system, we propose that the 1b and 1s inputs converge on each muscle and establish an activity set point that serves as a reference for synaptic perturbations. If this model is true, then upon loss of one input the muscle would sense a drop in presynaptic drive and instruct remaining MNs to compensate. Several retrograde pathways have been uncovered which lead to presynaptic homeostatic plasticity (Davis et al., 1998; DiAntonio et al., 1999; Haghighi et al., 2003; Hauswirth et al., 2018; Kauwe et al., 2016; Kikuma et al., 2019; Newman et al., 2017; Penney et al., 2012; Spring et al., 2016). Studies have found that decreased phosphorylation of CaMKII whether postsynaptic (Newman et al., 2017) or presynaptic (Li et al., 2018) is associated with induction of presynaptic homeostatic potentiation (PHP) in a synapse specific manner. The ablation of 1s MNs could be interpreted as a decrease in glutamate receptor function by the muscle and lead to compensatory changes at 1b MNs.

The above mechanism relies on the muscle recognizing that the 1s MN is missing. An alternative, unexplored model is that 1s MNs signal, either directly to 1b MNs or indirectly through the muscle, to coordinate postsynaptic activity. During larval development, co-innervating 1b and 1s NMJs are in very close proximity. Secreted or membrane-bound 1s signaling molecules could bind directly to receptors on 1b boutons to dampen 1b activity and establish the set point. Loss of the 1s input would release the inhibition and increase 1b-specific activity leading to functional compensation. Several studies have uncovered synaptic connectivity and development cues that are expressed in specific cells of the neuromuscular circuit, including Wnt4 (Inaki et al., 2007), Netrin B (Labrador et al., 2005; Mitchell et al., 1996; Winberg et al., 1998), Toll (Inaki et al., 2010; Rose et al., 1997), and Capricious (Kurusu et al., 2008; Nose, 2012; Shishido et al., 1998). In our previous work, we found *DIP-α* in two 1s MNs, and a *DIP-α* binding partner, Dpr6, in 1b and 1s MNs (Ashley et al., 2019). Additionally, other *DIP*s and Dprs are expressed in MNs (Carrillo et al., 2015). The potential combinatorial ligand-receptor interactions could contribute to the functional compensation at specific 1b inputs.

The neuromuscular circuit is a reductionist system with synaptic targets receiving up to four inputs. However, mechanisms uncovered at the NMJ can act in more complex circuits in vertebrates and invertebrates. For example, in the fly VNC, each MN receives sensory information from many interneurons to regulate motor behaviors (Heckscher et al., 2015; Kohsaka et al., 2019; Schneider-Mizell et al., 2016). The convergence of inputs on one MN may establish a homeostatic set point in the MN, similar to the muscle. The complex interneuron-MN connectivity more closely resembles that of vertebrate CNS neurons with polysynaptic dendrites (Heckscher et al., 2015; Kim et al., 2009; Kohsaka et al., 2019; Zarin et al., 2019). Additionally, age-(Bergado and Almaguer, 2002; Griffith et al., 2014; Mattson and Magnus, 2006; Mostany et al., 2013; Petralia et al., 2014) and disease-related (Gorman, 2008; Lepeta et al., 2016; Milnerwood and Raymond, 2010; Salvadores et al., 2017; Smith-Dijak et al., 2019) changes in synaptic function and cell survival have been observed in CNS and neuromuscular circuits. Thus, future studies at the NMJ and other circuits will elucidate the mechanisms governing how and when the activity set point is defined in a target-specific manner and how neurons respond to dysfunctional neighboring neurons.

### Update

While preparing this manuscript, the Littleton lab published a BioRxiv preprint that both confirms and complements our results (Aponte-Santiago et al., 2020). They utilized genetic ablations and manipulated neuronal activity to examine how 1b and 1s MNs respond to perturbations of adjacent inputs. They analyzed dorsal muscle 1, which is innervated by a different 1b MN and by the same 1s MN (dCE), and found similar results to ours: 1b NMJs expand and elevate activity upon loss of 1s inputs. In our study, we observed that 1b NMJs on ventral (m6 and m12) and dorsal (m4) muscles expanded upon loss of respective 1s inputs and increased their basal release probability, but compensation of evoked activity was different at each muscle, suggesting that input and/or muscle specific mechanisms modify each 1b synaptic terminal. Additionally, we found that the temporal dynamics of loss of 1s innervation are critical for enabling 1b structural and functional plasticity: co-innervation is required otherwise the 1b input does not respond. Together, both studies highlight that neurons have the capacity to alter synaptic terminal size and function when confronted with perturbations of converging inputs.

## Acknowledgements

We would like to thank Edwin “Chip” Ferguson, Ellie Heckscher, Kai Zinn, Kai Wang, Hong Liu, Richard Fehon, David Pincus, and members of the Carrillo lab for helpful discussions and comments. Stocks obtained from the Bloomington Drosophila Stock Center (NIH P40OD018537) were used in this study. The 3C10 anti-even skipped was deposited to the DSHB by Goodman, C. (DSHB Hybridoma Product 3C10 anti-even skipped), and was obtained from the Developmental Studies Hybridoma Bank, created by the NICHD of the NIH and maintained at The University of Iowa, Department of Biology, Iowa City, IA 52242. This work was supported by National Institutes of Health Grants K01 NS102342 (to RAC), T32 GM007183 (to ML-R), funding from the University of Chicago Biological Sciences Division (BSD) international student fellowship (to YW), funding from the BSD Office of Diversity and Inclusion (to RAC) and the Grossman Institute for Neuroscience, Quantitative Biology and Human Behavior (to RAC).

---

**Figure 2-1.**
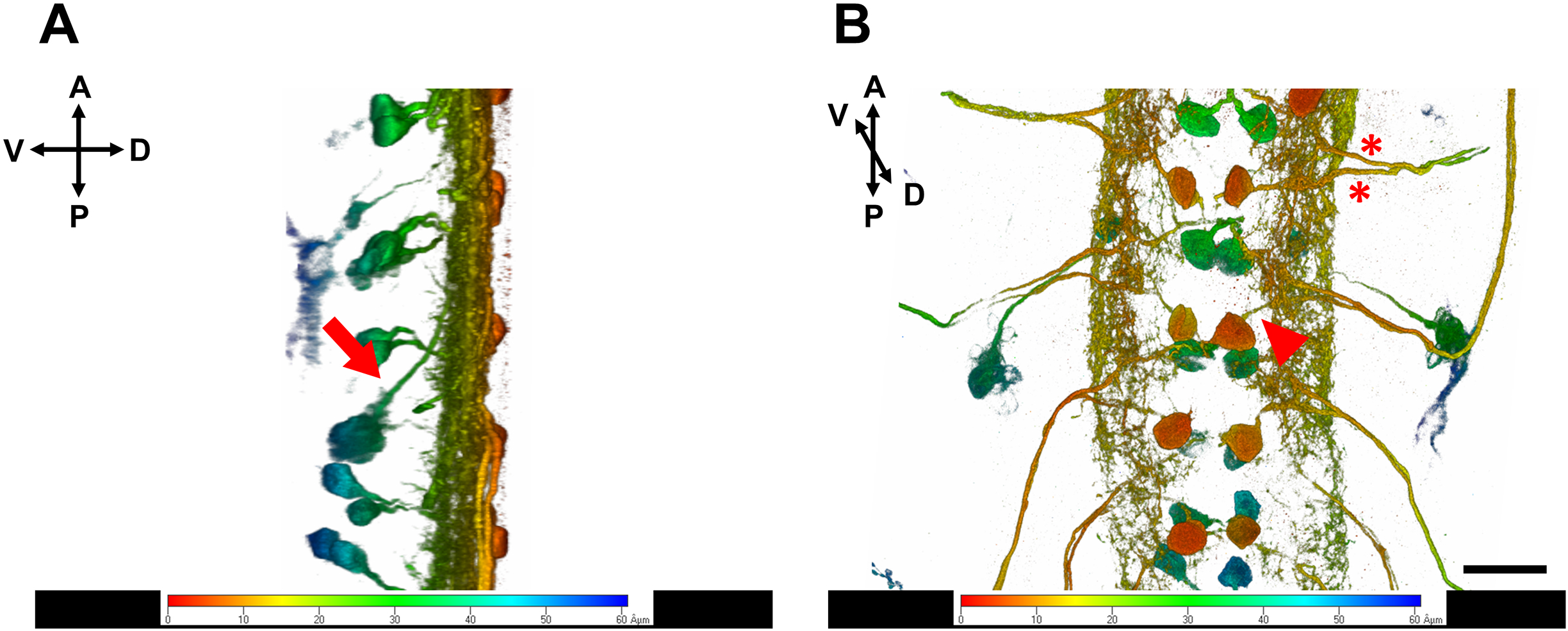
3D representations of A8 expressing neurons in the VNC. (**A,B**) Third instar larval VNC confocal stack projected in 3D and viewed from (**A**) lateral and (**B**) dorsal sides. Arrow in (**A**) shows a ventral cell (left, green) that projects an axon to the dorsal midline. Arrowhead in (**B**) shows an ipsilateral projection from a dorsal cell (orange). Red color denotes cells in the dorsal region of the VNC and green denotes deeper ventral cells. Asterisks indicate axons exiting the VNC. Calibration bar in is 22μm.

**Figure 6-1.**
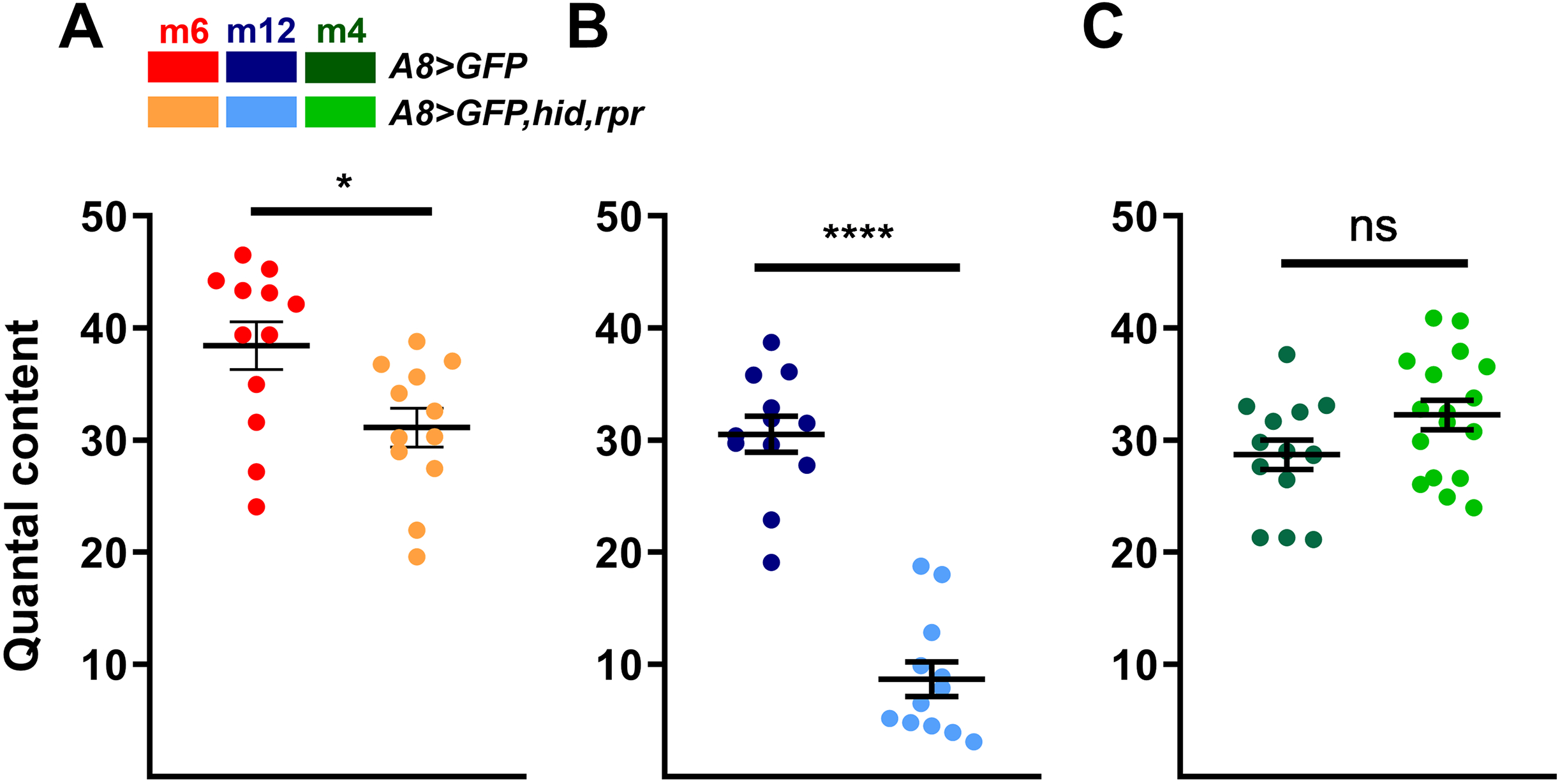
Quantal content is only compensated on m4 upon loss of 1s input. (**A-C**) Quantification of quantal content from (**A**) m6 (t(22)=2.664, p=0.0142, unpaired t test), (**B**) m12 (t(22)=9.885, p<0.0001, unpaired t test) and (**C**) m4 (t(29)=1.892, p=0.0685, unpaired t test) in control (A8>GFP) and mutant (A8>GFP,hid,rpr) animals. Quantal content was not decreased on m4, suggesting that m4-1b elevated vesicle release upon stimulation. Error bars indicate ±SEM. *p<0.05, ****p<0.0001. n (NMJs/larva) is 12/9, 12/11, 12/9, 12/12, 14/11, and 17/14 respectively.

**Figure 7-1.**
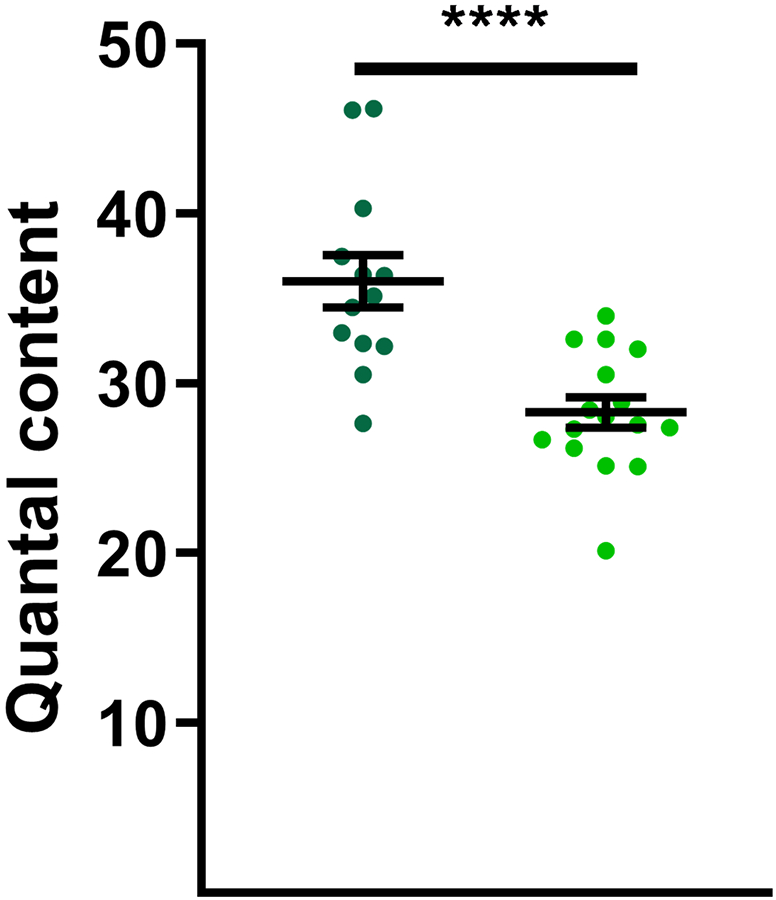
Quantification of m4 quantal content in control and mutant larvae (t(27)=4.572, p<0.0001, unpaired t test). Error bars indicate ±SEM. **p<0.01, ****p<0.0001. n (NMJs/larva) is 13/10 and 16/10 respectively.

